# Comparative genomic insights into bacterial induction of larval settlement and metamorphosis in the upside-down jellyfish *Cassiopea*

**DOI:** 10.1101/2022.06.24.497576

**Authors:** Aki Ohdera, Khushboo Attarwala, Victoria Wu, Rubain Henry, Henry Laird, Dietrich K. Hofmann, William K. Fitt, Mónica Medina

## Abstract

Bacterial biofilm is crucial in inducing the larval transition from pelagic to benthic environments for marine organisms. Bacteria can therefore dictate species distribution and success of the individual. Despite the importance of marine bacteria to animal ecology, the identity of inductive microbes for many invertebrates are unknown. We isolated bacteria belonging to multiple phyla are capable of inducing settlement and metamorphosis in the upside-down jellyfish *Cassiopea xamachana*. The most inductive isolates belonged to the genus *Pseudoalteromonas*, a marine bacterium known to induce the pelago-benthic transition in other marine invertebrates. In sequencing the genome of the isolated *Pseudoalteromonas* and an inductive *Vibrio*, we found biosynthetic pathways previously implicated in larval settlement were absent in these *Cassiopea* inducing taxa. Comparative analysis of the *Pseudoalteromonas* and *Vibrio* revealed shared genes that could underlie the inductive capacity of these two bacteria. Thus, *C. xamachana* are capable of responding to multiple bacterial species, but they may be responding to a common cue produced by multiple taxa. These findings could provide hints to the ecological success of *C. xamachana* compared to sympatric congeneric species within mangrove environments and provide avenues to investigate the evolution of animal-microbe interactions.

## Introduction

Microbe-animal interactions have shaped the evolution of metazoans, likely facilitating the emergence of multicellularity and the diversification of animal lineages (1, 2). In particular, larval settlement, the behavioral migration from the planktos to the benthos of some marine invertebrates, and metamorphosis, a developmental transition of life stage, is often triggered by microbe-animal interactions (3-6). Microbes thus dictate developmental timing of marine invertebrates, with implications for ecological and population dynamics of the animal. While abiotic factors including light, temperature, salinity, and substrate contribute to site selection for settlement, bacterial cues are found to be increasingly important to the life history of many marine animals (7-9).

Bacterial biofilms found on substrates are often sources of these cues, which can vary from water soluble compounds to physical cues (10-13). Detection of bacterial cues allow larvae to select appropriate habitats for subsequent adult survival, and can often be highly specific (14, 15), with larvae delaying settlement for prolonged periods when cues are undetected (16). Bacteria-associated site selection may also benefit the larvae, as some secondary metabolites exhibit anti-bacterial and anti-fungal properties, potentially acting as a “secondary immune system” for the animal (17, 18). Despite the ecological and evolutionary importance of biofilm to marine benthic communities and the growing number of microbial taxa identified to be settlement inducers, identification of the cues responsible for settlement and metamorphosis have been limited. Determining the diversity of cues and the sources of these cues is important to understanding the drivers that shape marine invertebrate life history.

In the upside-down jellyfish *Cassiopea xamachana*, microbial cues are predicted to initiate larval settlement and metamorphosis. Neumann (1979) showed *Cassiopea* metamorphose in response to the biofilm of *Vibrio* sp. under laboratory conditions. Cholera toxin isolated from *V. cholerae* was also found to induce settlement and metamorphosis (19), and an extract from collagen digested by *V. alginolyticus* was also found to be inductive (20). In their natural environment, brooded *C. xamachana* larvae are released into the water column and will often settle preferentially onto the underside of degrading mangrove leaves (7, 21). When treated with antibiotics, larval settlement and metamorphosis is abolished in response to mangrove leaves. These findings suggest bacteria found in biofilm associated with degrading leaves may be a source of the inductive cue of *Cassiopea*. In a follow-up study, Fleck and Fitt (7) showed a water soluble, 5.8 kDa proline-rich peptide extracted from degraded mangrove leaves to be a an inducer of settlement in *C. xamachana*. However, the originating bacterial source of the cue, and whether the cue is a byproduct of leaf degradation, remains unknown.

A large body of work has offered clues (14, 15) for how settlement and metamorphosis occurs through animal-microbe interactions. Yet, we do not fully understand the extent of the specificity underlying the developmental transition of pelagic larvae to benthic juveniles. Here, we investigate substrate preference of two *Cassiopea* species found to occur sympatrically in a mangrove environment. We isolated bacterial species from select substrate to identify bacteria capable of inducing the pelago-benthic transition in *C xamachana*. Comparative genomics of bacterial isolates capable of inducing settlement and metamorphosis revealed potential mechanisms underlying induction in *C. xamachana*. Determining larval settlement in *Cassiopea* can further our understanding of animal-microbe interaction dictating development. In addition, the development of the jellyfish as a model to study host-microbe interactions necessitates identification of bacterial inducers of larval settlement.

## Results

### Settlement of *C. xamachana* on Natural Substrates

As with previous studies (7, 21), *C. xamachana* settled and metamorphosed in response to natural substrates. We found larvae responded to both organic and inorganic substrates, including sand and bivalve shell fragments collected from onshore and offshore sites (Fig. 1A, B). Response to inorganic substrate (sand, bivalve shell) may indicate that an inductive cue is directly produced by biofilm associated bacteria, and is not a plant degradation product. When compared to *Cassiopea frondosa*, larvae of *C. xamachana* appeared to be generalists, settling onto multiple substrate types (Fig. 1). Interestingly, mangrove leaves, a known inductive substrate, exhibited the lowest settlement compared to onshore sand in *C. xamachana*. On the other hand, *C. frondosa* exhibited a weak response to the all substrates. These results suggest either settlement and metamorphosis inducing taxa of *C. xamachana* are common across substrate types, or *C. xamachana* responds to a wide range of cues. Unlike *C. xamachana*, inductive taxa of *C. frondosa* are rare and larvae respond specifically to one or a few cues.

**Figure 1:**
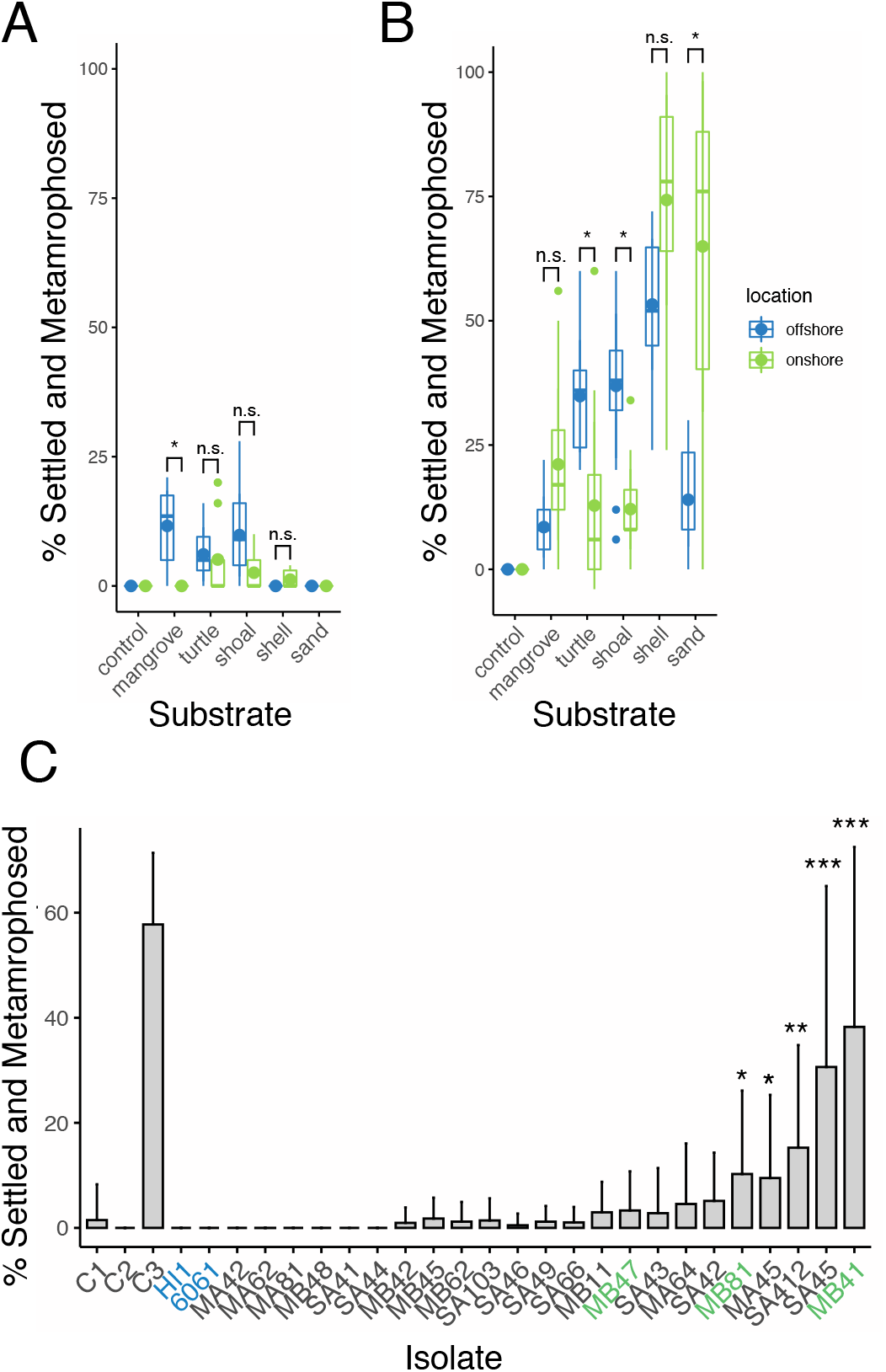
Settlement and metamorphosis of *Cassiopea* spp. in response to natural substrate and monoculture bacterial biofilm. A) Settlement of *Cassiopea frondosa* in response to natural substrate. Settlement and metamorphosis was monitored over the course of four days. B) Settlement of *Cassiopea xamachana* in response to natural substrate. C) Settlement and metamorphosis of *C. xamachana* larvae in response to monoculture biofilm of bacteria isolated from degrading mangrove leaves and sand from onshore sites. *Pseudoalteromonas* isolates are shown in green. *Pseudoalteromonas luteoviolacea* species known to induce other invertebrate taxa are shown in blue. Statistically significant rates of settlement and metamorphosis are indicated. * = < 0.05, ** < 0.005, *** < 0.0005 (Kruskal-Wallis, Wilcoxon test). Settlement rates were compared to controls. C1= 0.45 µM filtered seawater. C2 = Marine broth pre-treatment C3 = Inductive peptide z-GPGGPA.

In order to determine whether microbes associated with inductive substrates can induce settlement and metamorphosis in *C. xamachana*, we isolated bacteria from mangrove leaves and sand. Settlement rate of *C. xamachana* larvae exposed to monoculture biofilms of the bacterial isolates varied (Fig. 1C). Larval metamorphic induction rates ranged from 0 % to nearly 40 % over 2 days. In particular, isolate MB41 and SA45 exhibited the greatest metamorphic induction of the tested isolates, with an induction rate of 38.25 % and 32.07 %, respectively. All other isolates either failed to induce settlement or did not exceed 10 % settlement on average. Altogether, five of the isolates elicited a significant settlement and metamorphosis response in *C. xamachana* larvae (Kruskal-Wallis, *post hoc* Wilcoxon test, p-value < 0.05) (Fig. 1C).

Broad classification using the 16S rRNA gene revealed the inductive isolates belonged to diverse bacterial groups (Table S1). The five most highly inductive taxa were identified as belonging to Pseudoalteromonadaceae, Sphingomonadaceae, and Rhodobacteraceae. In particular, isolates MB41, MB47, and MB81 belonged to genus *Pseudoalteromonas*, closely related to *Pseudoalteromonas lipolytica* and *P. donghaensis* (Fig. 2). Given that *Pseudoalteromonas luteoviolacea* is a well characterized settlement inducer of the polychaete worm *Hydroides elegans* (10, 22, 23), we tested whether strains of *P. luteoviolacea*, induced settlement in *C. xamachana* larvae. Monoculture biofilm of *P. luteoviolacea* HI1 and 6061 failed to induce settlement and metamorphosis, and were lethal after 48 hr. Another isolate of interest, although not a significant inducer, MA64 was identified as closely related to *Vibrio alginolyticus*. In prior studies a *Vibrio* was found to induce settlement in *C. andromeda* (24, 25).

**Figure 2:**
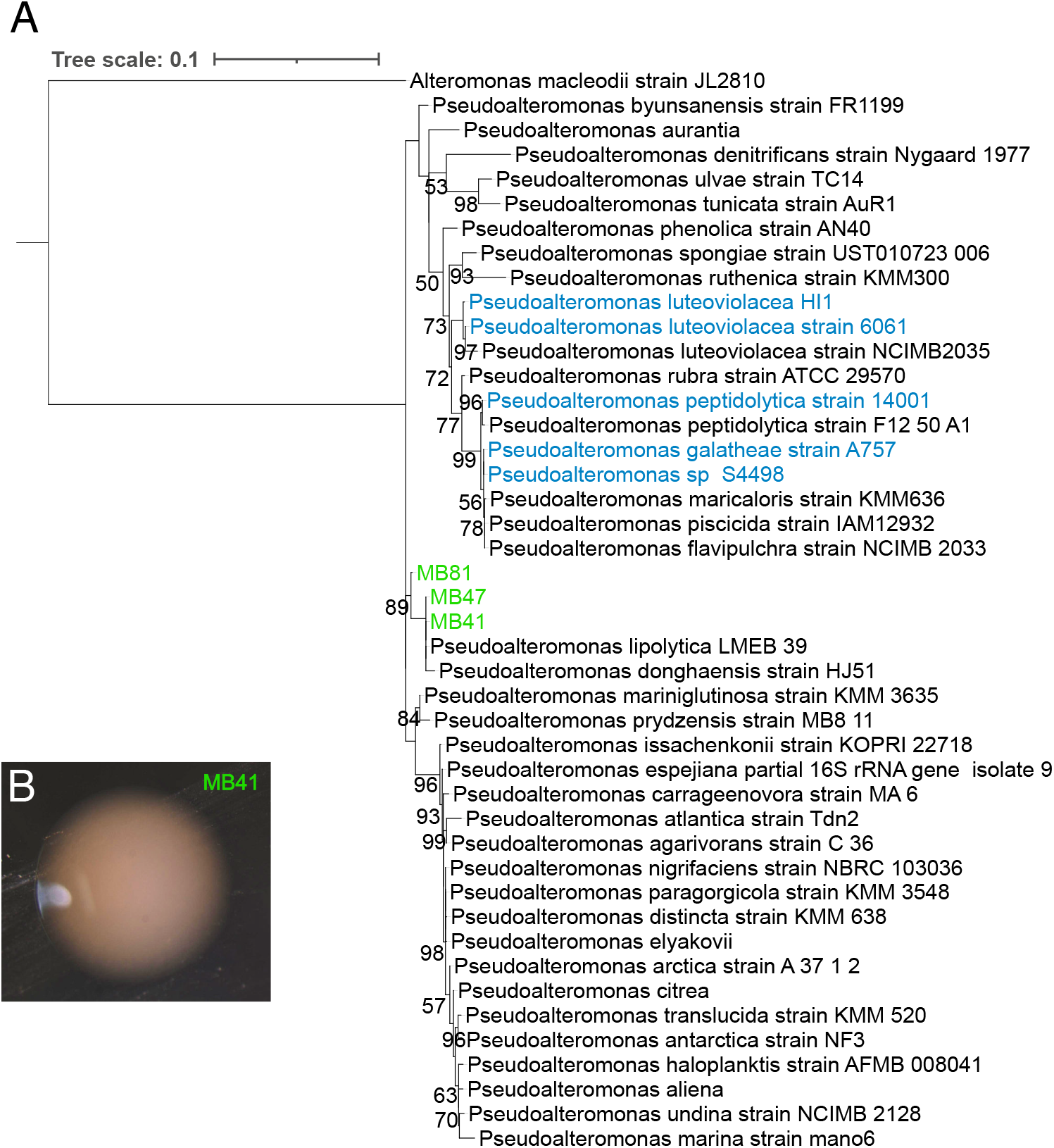
Phylogenetic reconstruction of genus *Pseudoalteromonas* using the 16S rRNA genes. Isolates are indicated in green. *Pseudoalteromonas* species used in the analyses are indicated in blue. Sequences were aligned using MAFFT, with trimming performed with BMGE. Construction of the phylogenetic tree was performed with iqTree. B) A colony on marine agar of *Pseudoalteromonas* sp. MB41.

### Inductive isolates are rare in degrading mangrove leaves

To determine whether inductive taxa were found to be common in substrate-associated biofilm, we performed amplicon sequencing of degrading mangrove leaves from three onshore sites and one offshore site. We found the microbiome of mangroves leaves to be highly diverse, and identified approximately 3000 ASVs with greater than 5 reads across the samples. The most abundant groups comprised roughly 25% of the total reads per sample. The five most common family of bacteria associated with mangroves leaves were Vibrionaceae, Sandaracinaceae, Rhodobacteraceae, Saprospirae, and Spirochaetaceae. Community composition differed significantly between several sites (Permanova, 9999 permutations, q-value < 0.05), including between the offshore site and an onshore site (Fig. 3, Fig. S1A). However, community composition was not significant when Benjamini-Hochberg FDR correction was applied for pairwise comparisons between sites (Fig. S1B).

**Figure 3.**
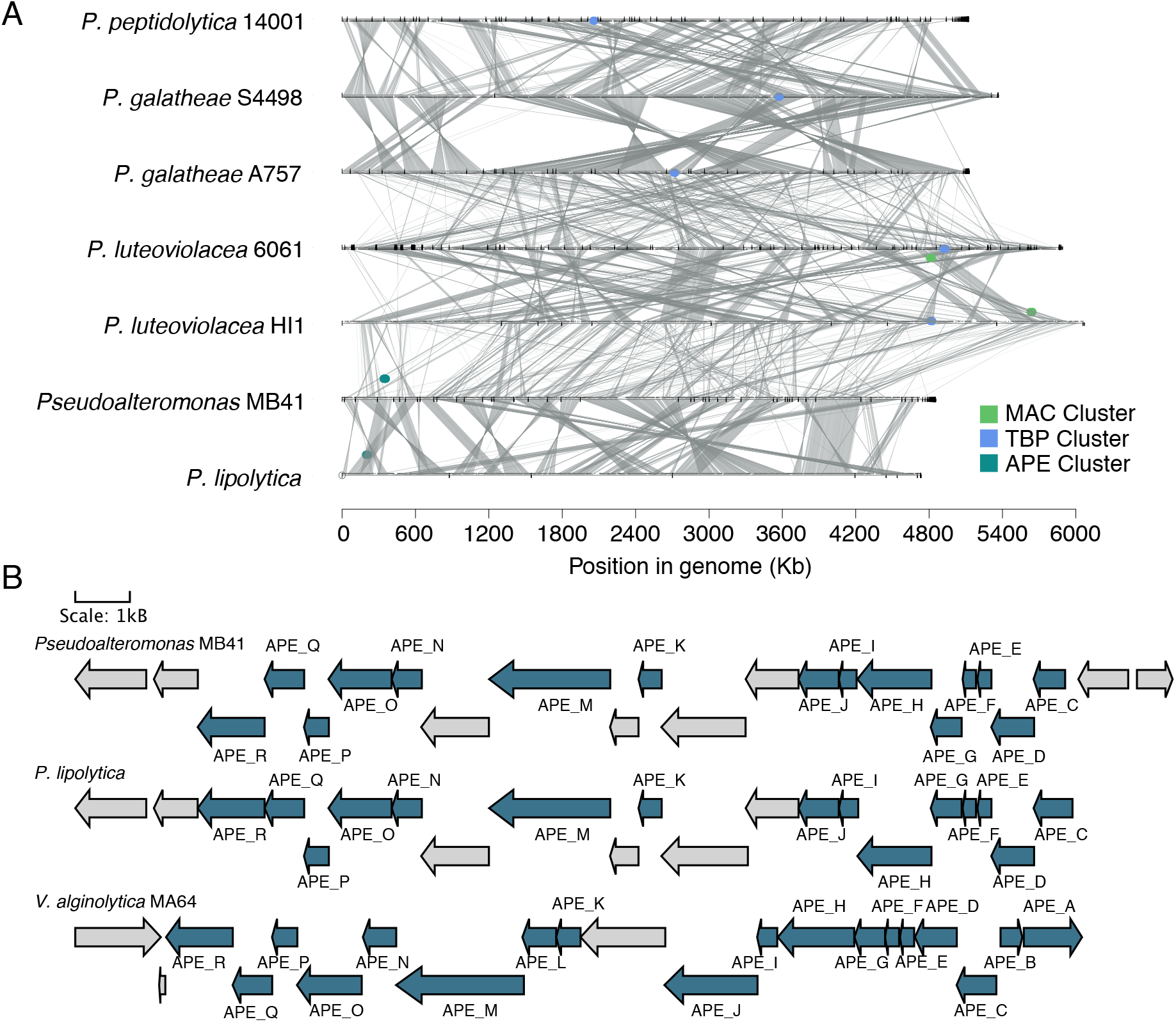
Heatmap of the top 25 most abundant family of bacteria identified from mangrove leaves collected from one offshore site and three onshore sites. Taxonomic identity and abundance (log10 transformed) were determined with Qiime2 against the Silva database.

Using a predefined cutoff, we identified ASVs matching the V4 region of the 16S rRNA gene of the bacterial isolates. We found that inductive isolates were rare within the mangrove microbiome for all sites (Fig. 3). In general, inductive taxa composed less than 1% of the total reads generated for each sample. Of the inductive isolates including those that elicit low settlement rates in *C. xamachana* larvae, a *Vibrio* sp. was the only ASV present that comprised greater than 1 % of total reads in the mangrove leaf microbiome.

### Genome Analysis of Inductive Isolates

The genus *Pseudoalteromonas* have been implicated in larval settlement and metamorphosis across many marine species (3, 26, 27). We therefore sequenced the genome of the highly inductive *Pseudoalteromonas* isolate MB41 to identify potential genomic signatures associated with induction of larval settlement. We generated a 4.85 Mb assembly with 4452 predicted coding genes. Average nucleotide identity (ANI) was greater than 98% when compared with *Pseudoalteromonas lipolytica* strain CSB02KR. Our genome assembly was screened for characterized settlement inducers of other *Pseudoalteromonas* spp.

*Pseudoalteromonas luteoviolacea* produces tailocins, a well characterized inductive cue of settlement and metamorphosis of *Hydroides elegans* (10, 11, 28). Composed of viral sheath components, tailocins are encoded the *MAC* protein gene cluster (Huang et al. 2012). In our isolate MB41, the tailocin gene cluster was absent as well as in the closely related *P. lipolytica*, revealing that the inductive cue of MB41 is not tailocins (Fig. 4A, Fig. S2).

**Figure 4:**
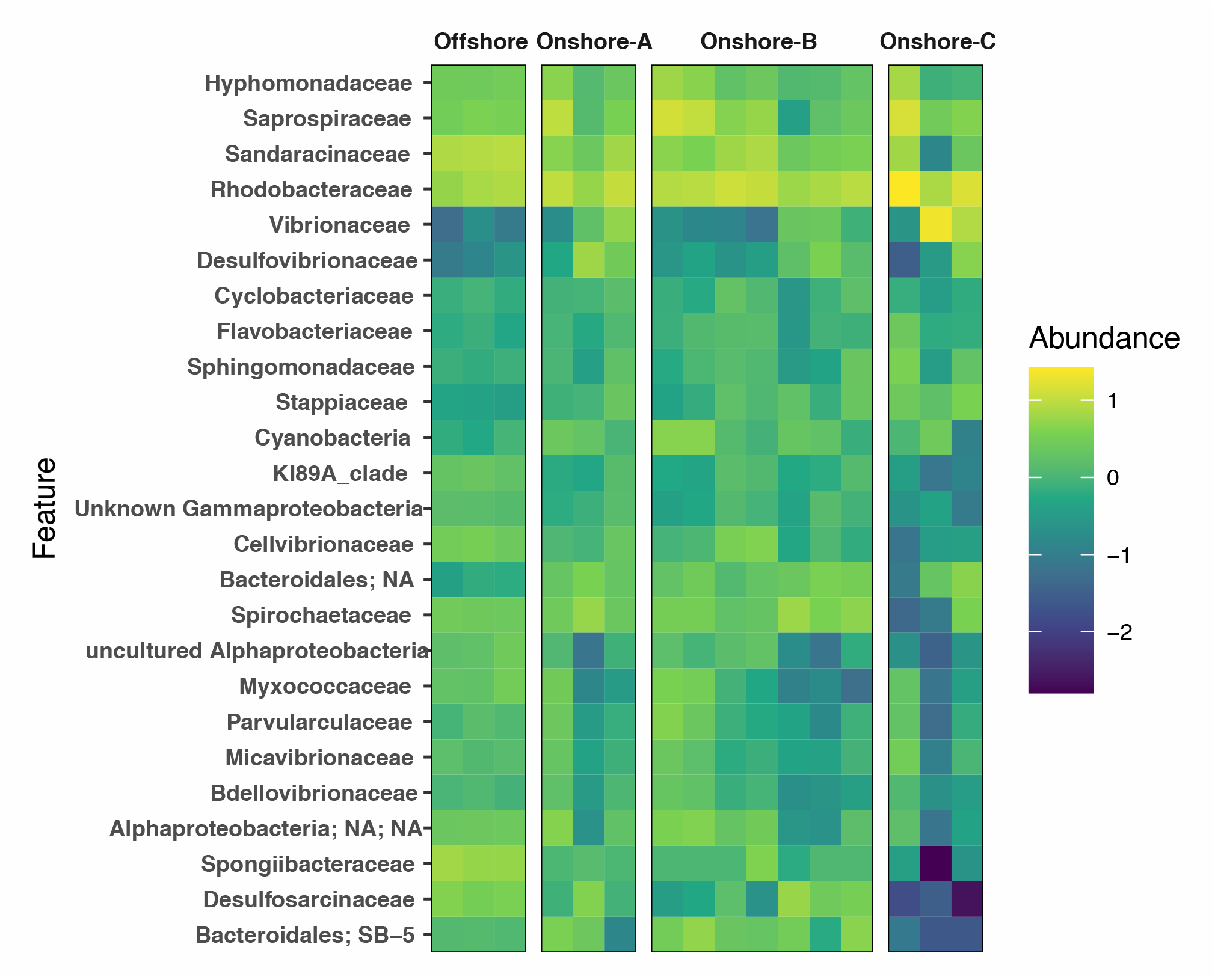
Comparative genomics of *Pseudoalteromonas* spp. A) Genome synteny of *Pseudoalteromonas* spp. Colored points indicate genomic position of biosynethic gene clusters. MAC = Tailocins. TBP = Tetrabromopyrrole. APE = Aryl polyene. B) Biosynthetic gene cluster of aryl polyene found in *Pseudoalteromonas* MB41, *Pseudoalteromonas lipolytica*, and *Vibrio alginolyticus* MA64. Genes previously confirmed to be involved in biosynthesis are colored in blue.

In addition to tailocins, the *bmp*1-10 biosynthetic gene cluster (BGC) of *Pseudoalteromonas* encodes for a set of enzymes known to produce the brominated compound tetrabromopyrrole. Tetrabromopyrrole was first identified to induce settlement and metamorphosis of corals (13, 29). Species in our analysis known to produce tetrabromopyrrole possessed a minimum of 6 of the 10 genes within the BGC, while all *bmp* genes were absent from the *P. lipolytica* and MB41 genomes (Fig. 4A). We therefore predicted additional BGCs with antiSMASH (30). Unlike the genomes of the pigmented *Pseudoalteromonas*, which includes *P. luteoviolacea*, few BGCs were predicted from our isolate MB41. Only two putative BGCs, an aryl polyene (APE) and ribosomally-synthesized and post-translationally modified peptide (RiPP) biosynthesis clusters were identified from MB41. APEs are polyunsaturated carboxylic acids, often found in the outer membranes of host-associated bacteria. BLAST alignments of the MB41 APE genes against previously described clusters suggested that the MB41 cluster is most similar to the flexirubin-like APE cluster found in *Vibrio fischeri* (31), although the biosynthetic product of the MB41 cluster remains unknown (Table S2). Of the *Pseudoalteromonas* genomes surveyed, APE clusters were only found in MB41 and *P. lipolytica* (Fig. 4). The RiPP cluster in MB41 was characterized by a DUF692, previously linked to the production of bacteriocins, or antibacterial peptides (32). Although the exact product from the Rip clusters are unknown, we found these clusters to also be present in both the *Pseudoalteromonas* HI1 and *P. lipolytica* genomes (Table S3).

As the most abundant inductive genus, albeit not the highest inducer, we also sequenced the genome of the *Vibrio* isolate MA64 producing a 5.5 Mb assembly with 5256 coding sequences. MA64 was highly similar to *V. alginolyticus* strain 12G01, with an ANI value greater than 98%. Similar to *Pseudoalteromonas* isolate MB41, the MA64 genome lacked both tailocin and tetrabromopyrrole BGCs. Secondary metabolite cluster prediction resulted in six putative clusters, including a putative APE (Fig. 4B) and RiPP cluster. In addition, we identified a cluster most similar to vibrioferrin, ectoine, and betalactone, as well as a 100% match to the vanchrobactin BGC. Like *Pseudoalteromonas* isolate MB41, the MA64 APE cluster was most similar to flexirubin-like cluster found in *V. fischeri* (Fig. 4B). The RiPP cluster of MA64 was also characterized by a DUF2063 and likely the core peptide.

We also searched for genes unique to *Pseudoalteromonas* isolate MB41 and *Vibrio* isolate MA64 by identifying orthologous groups, hereafter called orthogroups. We identified 26 orthogroups with genes only shared between MB41 and MA64 (Supplemental File 1). Of note, genes with dynamin-like domains were shared between MB41 and MA64 but absent from all other genomes analyzed, including *P. lipolytica*. Bacterial dynamin-like genes are implicated in extracellular vesicle formation and a wide range of functions involving membrane dynamics. Interestingly, the dynamin-like genes found in *Pseudoalteromonas* isolate MB41 were most closely related to proteins found in *Vibrio* spp. In *Pseudoalteromonas* MB41, the dynamin-like domain proteins are found in tandem, while a single gene was found in *Vibrio*.

## Discussion

Bacteria have been shown to be a source of larval settlement cues for marine invertebrate species, but taxa capable of inducing settlement and metamorphosis of larvae remains largely unknown (25, 29, 33-37). We found, larvae of *C. xamachana* appear to be broad in specificity, with bacteria from multiple phyla inducing the pelago-benthic transition in larvae (Fig. 1). Two of the inductive isolates were identified as belonging to the marine genus *Pseudoalteromonas* (MB41) and the nonpathogenic *Vibrio* closely related to *V. alginolyticus* (MA64). The latter has previously been shown to induce settlement and metamorphosis in *C. andromeda* but this is the first record for a jellyfish settlement inducing *Vibrio* species isolated from a natural substrate. Given the ubiquity of *Pseudoalteromonas* in the literature as an inducer of larval metamorphosis (El Gamal et al. 2016; Huang et al. 2007), the discovery of a *Pseudoalteromonas* species capable of inducing metamorphosis of a scyphozoan (MB41) while another is lethal (*P. luteoviolacea)* highlights the diverse role the group may play in the ecology of marine invertebrates.

Interestingly, inductive bacteria were rare within the microbial community of degrading mangrove leaves (Fig. 3). Similar observations have been made previously and may suggest an underappreciated specificity to biofilm associated bacteria by larvae (38, 39). In fact, some animal species preferentially settle in proximity to rare microbial taxa (40). Larvae may also respond to multiple taxa or multiple cues, including bacterial species that may be difficult to culture but common within the biofilm. This confounds the exact animal-microbe interaction that drives the pelago-benthic transition for *Cassiopea*. However, given the difference in settlement rate between onshore and offshore sites for both *Cassiopea* species, bacterial community composition likely remains an important factor in dictating the larval transition. Furthermore, the behavior in *C. xamachana* suggests redundancy in cues, which can be predicted to have ecological and evolutionary implications. For example, *C. frondosa* occur at lower rates in the Florida Keys compared to *C. xamachana (41)*. While several factors may contribute to this difference in species distribution, the biofilm composition on available substrate may play an important role in regulating abundance of the two *Cassiopea* species within the environment of our study. In addition, our data suggest a plant substrate is unnecessary for settlement of *C. xamachana* larvae and inductive bacteria do not require exogenous molecules as precursors to produce the settlement inducing molecule(s). These findings thus indicate the importance of bacteria for larval settlement of *Cassiopea*.

In searching for inductive biosynthetic gene clusters (BGCs), we found the *Pseudoalteromonas* isolate MB41 and *Vibrio* isolate MA64 genomes both lack genes responsible for tetrabromopyrrole biosynthesis, as well as tailocins in the case of *Pseudoalteromonas*, although the presence of the tailocin genes cluster is not predictive of settlement induction (42) (Fig. 4). Similar to previous findings, tailocin gene clusters were only found in *P. luteoviolacea*. The *bmp* gene cluster responsible for tetrabromopyrrole production typically contains 10 genes and is commonly found in pigmented *Pseudoalteromonas* species. The *bmp* gene cluster was absent from the genome of the unpigmented *Pseudoalteromonas* isolate MB41. These findings suggest *Cassiopea* settle and metamorphose in response to cues yet to be identified from a *Pseudoalteromonas*.

Another hypothesized route of inductive cue delivery/release is through outer membrane vesicles (OMVs). OMVs produced by *Cellulophaga lytica* carrying lipopolysaccharides were shown to induce settlement and metamorphosis in *H. elegans* (43). OMVs are also known to carry various payloads, including toxins, genetic material, lipids, etc., and may be a common mechanism through which larvae and biofilm associated microbes interact (43). OMVs have been identified from several *Pseudoalteromonas* and *Vibrio* species, and may be involved in inducing settlement in *Cassiopea* (Nevot et al. 2006, Bitar et al. 2019). A unique dynamin-like protein was identified from the genomes of *Pseudoalteromonas* isolate MB41 and *Vibrio* isolate MA64, which have been associated with the production of extracellular OMVs (44, 45). These may be indicative of similar processes leading to larval settlement and metamorphosis in *Cassiopea*, but additional experiments will be necessary to confirm this.

We also identified putative clusters responsible for aryl polyenes clusters (APE) biosynthesis. APEs are polyketide derivatives, a class of pigments related to carotenoids and common in gram-negative bacteria (46). Neither *Pseudoalteromonas* isolate MB41 and *Vibrio* isolate MA64 are considered pigmented, yet possess the BGCs (Fig. 4). MB41 belongs to the non-pigmented group within the *Pseudoalteromonas* genus (Fig. 2). Loss of pigmentation has been associated with reduction in antifouling activity of mutant strains of pigmented species (47, 48). Of the four classes of aryl polyenes, the cluster found in MB41 and MA64 were most similar to flexirubin-like BGC found in *Vibrio fischeri* (Cimermancic 2014). As with other APEs, the flexirubin-like APE is thought to be antioxidative, but further investigation will be required to determine its involvement as an inductive metabolite of invertebrate settlement and metamorphosis.

Despite the importance of bacterial-larval interactions to animal evolution and ecology, we know little of the inductive cues responsible for larval settlement and metamorphosis. Given the wealth of information available through genomic data, understanding genomic features underlying settlement and metamorphosis can accelerate discovery of additional mechanisms driving the developmental transition. Understanding larval settlement provides information on species distribution and can aid in restoration efforts, particularly in the case of ecologically important scleractinian coral species (49).

## Materials and Methods

### Larval Collection

Brooding females of *C. xamachana* were collected from a sea grass bed off of Key Largo, FL (25.102204, -80.438708, 24.749522, -80.978341) (Fig. S1A). Female *C. frondosa* were collected from (25.128443, -80.443558). Eggs were removed from the females by gently pipetting seawater at the central brood vesicles and released embryos were collected into 0.22 µM filtered artificial sea water. Developing embryos were transferred to glass finger bowls containing antibiotic seawater (100 µg/ml neomycin, 130 µg/ml streptomycin). Embryos were left undisturbed for 48 hrs to allow development into mobile planulae. The planulae were transferred to fresh antibiotic seawater (ABS) and used for settlement assays within 24 hr.

### Bacterial Isolation

Degraded mangrove leaves were collected from below mangrove stands and rinsed gently three times with 0.2 µm filtered seawater (FSW). 1 cm^2^ pieces were cut from the leaves and homogenized in 25 ml of FSW using an ethanol sterilized mortar and pestle. Serial dilutions of the homogenates were made by transferring 0.5 ml of the diluted homogenate into 9.5 ml of FSW for 5 dilutions. 100 µl of each dilution was plated onto marine agar (Difco Marine Agar 2216) and incubated at 28°C for 24 hr. Bacterial colonies were picked and preserved in FSW with 15% glycerol at -80°C.

### Identification of Bacterial Isolates

The full length of the 16S ribosomal RNA gene of the bacterial isolates was sequenced using universal primer pair 27F and 1492R (50). Initially, bacterial stocks were plated onto marine agar and grown for 48 hrs. Colonies were picked and added directly to the PCR mix and the 16S gene was amplified using the following cycling conditions: 95 for 5 min, 94 for 1, 55 for 1:30, 72 for 2:00, 72 for 5:00, for 35 cycles. Sanger sequencing of amplicon products was performed at the Pennsylvania State University Sequencing Core Facility. Chromatograms were manually curated using MEGA, and sequences were compared to both RDP and NCBI databases for taxonomic identification. Phylogenetic analysis of isolates belonging to the genera *Pseudoalteromonas* and *Vibrio* was performed as follows: First the 16S rRNA gene sequences were aligned with MAFFT (ver. 7.271). Alignments were trimmed with BMGE (ver. 1.12). IQ-TREE (ver. 1.6.12) was used for phylogenetic inference, with model selection performed with ModelFinder (Kalyaanamoorthy et al. 2017). Tree visualization and curation was performed with iTOL (ver. 6).

### Settlement Assays

Bioassays were prepared in 12 well plates by filling individual wells with 1 ml of sterile marine media (Difco Marine Broth 2216). Control wells were filled with filtered seawater (FSW). A second negative control was included in which wells pre-treated with marine media were left un-inoculated and rinsed with FSW as described below. Wells containing media were inoculated with isolates by transferring a single colony to a well. Wells were inoculated in triplicate per isolate, and were randomly assigned across multiple plates. Plates were incubated at 28°C for 24 hrs without shaking to allow the bacteria to grow and settle on the surfaces of the well. Wells were gently rinsed with 1 ml of FSW without disrupting the biofilm for a total of 2 washes at the end of the 24 hrs. Wells that were still turbid after the two washes were rinsed for a third time, or until the seawater remained clear. Wells were then filled with a total of 2 ml of 0.2 µm FSW. A positive control was included in the assay where the synthetic settlement and metamorphosis inducing peptide (GPGGPA) were added at a final concentration of 1.5 × 10^−5^ M. Approximately ten *C. xamachana* larvae kept in ABS for 48 hrs were placed in each of the wells. Plates were kept undisturbed at 25°C in the dark. The plates were scored for settlement and metamorphosis at 24 and 48 hrs using a Stemi 305 stereo microscope (Zeiss).

### 16S microbiome extraction and sequencing

Degraded mangrove leaves were collected during the summers of 2013 and 2015 in Key Largo Florida from five different locations (Figure 1). Leaves were immediately stored at -20°C until further processing. DNA was extracted using the PowerSoil DNA Isolation Kit (Mobio) following the manufacturer protocol. The V4 region of the 16S rRNA gene was PCR amplified using the 515F (GTGCCAGCMGCCGCGCGGTAA) -806R (GGACTACHVGGGTWTCTAAT) primer pair (51). Sequencing was performed at the Joint Genome Institute (JGI) on an Illumina MiSeq with 300 bp paired-end run mode.

Reads were analyzed using Qiime2 (ver. 2021.4). Reads were initially processed by JGI according to their iTag analysis protocol, followed by additional processing and analysis. Briefly, read trimming, filtering, and denoising were performed with the Dada2 plugin (Callahan et al. 2016). Taxonomic assignment was performed with ‘feature-classifier classify-sklearn’, using the Naïve Bayes classifier trained on the Silva_138_99%_505F/806R region specific database (Bokulich et al.) Visualizations were performed with Qiime2R (ver. 0.99.6).

### Genome Sequencing and Analysis

Bacterial isolates MB41 and MA64 were chosen for whole genome sequencing based on their inductive capacities. Colonies were grown in marine media for 24 hrs. Genomic DNA was extracted following the JGI Bacterial DNA Isolation CTAB-2012 protocol (https://jgi.doe.gov/user-program-info/pmo-overview/). Illumina sequencing libraries were constructed using the Illumina TruSeq Nano DNA prep Kit with a 550 bp insert size. Libraries were sequenced on the Illumina Miseq, generating 2.8 and 2.7 Gb of 300 bp paired-end sequence data for MB41 and MA64, respectively. Draft genomes were *de novo* assembled using the A5-miseq pipeline (52). Genomes of *Pseudoalteromonas luteoviolacea* strains HI1 (53) and 6061 (54) were downloaded from NCBI (accession JWIC01000000, CAPN00000000). Genes were annotated with Glimmer3.0 implemented through RAST (55). The presence of tailocin encoding *mac* protein genes, tetrabromopyrrole gene cluster, and aryl polyene clusters were determined using BLAST 2.5.0+ (56). Secondary metabolite cluster prediction was performed with antiSMASH (ver. 6.0) (Blin et al. 2019). Visualizations were performed with Gene Graphics (Harrison et al. 2017) and Synima (Farrer 2017).

## Data Availability

Raw read and genomic assemblies of *Pseudoalteromonas* MB41 and *Vibrio* MA64 have been deposited in the NCBI/GenBank database under the project accession PRJNA774281.

## Acknowledgments

We are grateful for the input and training on experimental methods bacterial isolation provided by Dr. Kristen Marhaver, as well as edits provided by Dr. Sheila Kitchen. Funding for this work was provided by NSF (OCE 1442206) and the Department of Energy (CSP 1622).

## Supplemental Figures and Tables

**Figure S1:**
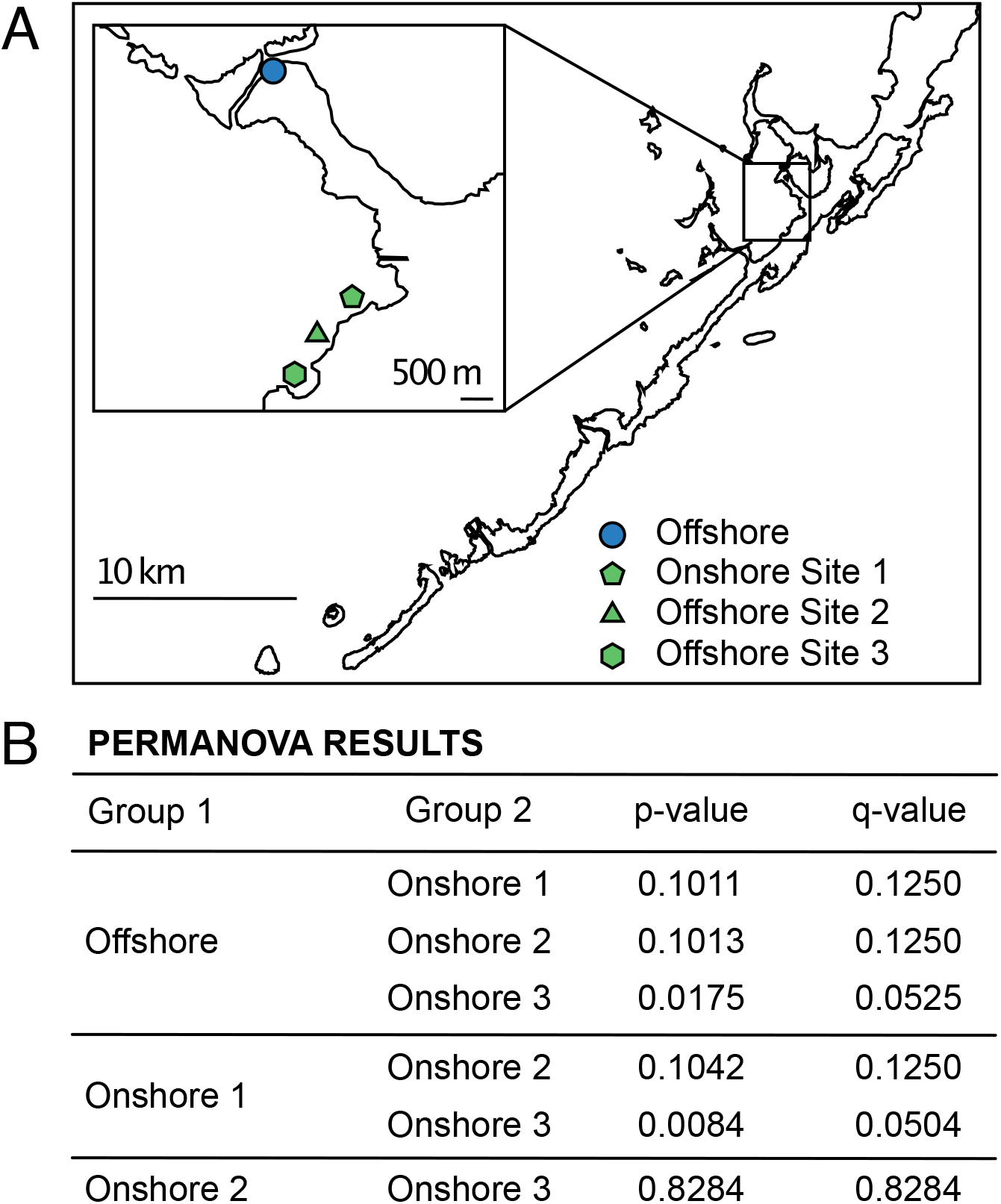
Onshore and offshore collection sites of degrading mangrove leaves in the Upper Florida Keys (A). Natural substrates were collected from offshore (blue circle) and onshore sites (green shapes). Results of permanova analysis of the microbial community associated with degrading mangrove leaves (B). Permanova analysis was performed with Qiime2.

**Figure S2:**
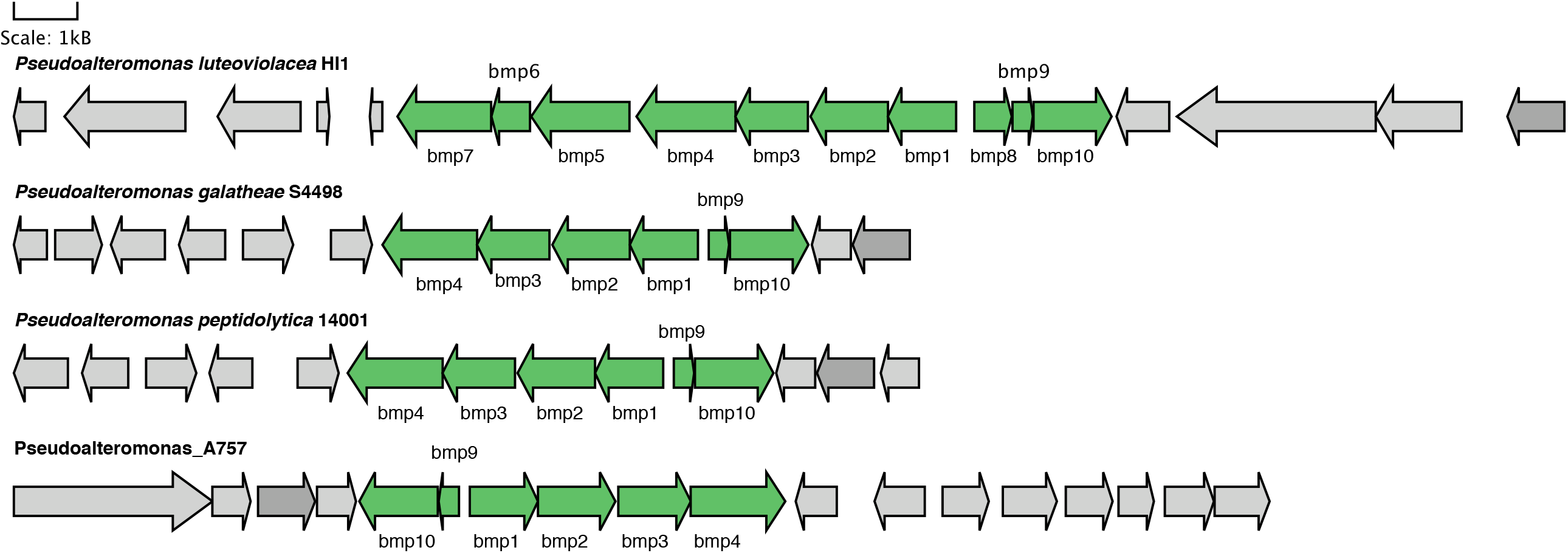
Tetrabromopyrrole synthesizing *bmp* gene cluster identified from pigmented *Pseudoalteromonas* species.

**Table S1.** Bacterial isolates collected from highly degraded mangrove leaves. Taxonomic placement of the isolates was performed through the analysis of full length 16S genes against the NCBI database.

| Isolate ID | Substrate | Induction | Taxonomic ID |
| --- | --- | --- | --- |
| MA42 | ML | N.I. | *Halomonas* sp. |
| MA45 | ML | Ind | Unknown |
| MA62 | ML | N.I. | *Thalassospira* sp. |
| MA64 | ML | Ind | *Vibrio* sp. |
| MA81 | ML | N.I. | *Pseudoalteromonas* sp. |
| MB11 | ML | Ind | *Pseudomonas* sp. |
| MB41 | ML | Ind. | *Pseudoalteromonas* sp. |
| MB42 | ML | Ind. | *Agarivorans* sp. |
| MB45 | ML | Ind. | Unknown |
| MB47 | ML | Ind. | *Pseudoalteromonas* sp. |
| MB48 | ML | N.I. | Unknown |
| MB62 | Sand | Ind. | *Gallaecimonas* sp. |
| MB81 | Sand | Ind. | *Pseudoalteromonas* sp. |
| SA41 | Sand | N.I. | *Erythrobacter* sp. |
| SA42 | Sand | Ind. | Unknown |
| SA43 | Sand | Ind. | Unknown |
| SA44 | Sand | N.I. | *Halobacillus dabaensis* |
| SA45 | Sand | Ind. | *Paracoccus* sp. |
| SA46 | Sand | Ind. | *Exiguobacterium* sp. |
| SA49 | Sand | Ind. | Unknown |
| SA412 | Sand | Ind. | *Erythrobacter* sp. |
| SA66 | Sand | Ind. | *Halomonas* sp. |
| SA103 | Sand | Ind. | *Nitratireductor* sp. |

\* N.I. = Non-inductive
\*\* Ind = Inductive

**Table S2.** Genes that comprise the aryl polyene (APE) cluster identified in *Pseudoalteromonas* sp. MB41, *Pseudoalteromonas lipolytica*, and *Vibrio* sp. MA64. Presence of APE genes were determined by antiSMASH prediction and confirmed by BLAST against the APE cluster identified in *E. coli*, where the IDs are borrowed from (Johnston et al. 2021).

|  | MB41 | MA64 | P. lipolytica | Gene Name |
| --- | --- | --- | --- | --- |
| APE_R | 6666666.166623.peg.884 | 6666666.64559.peg.2282 | gene-AT00_RS12545 | Putative beta-ketoacyl-ACP synthase |
| APE_Q | 6666666.166623.peg.885 | 6666666.64559.peg.2283 | gene-AT00_RS12550 | 3-oxoacyl-[acyl-carrier protein] reductase |
| APE_P | 6666666.166623.peg.886 | 6666666.64559.peg.2284 | gene-AT00_RS12555 | Conserved hypothetical protein |
| APE_O | 6666666.166623.peg.887 | 6666666.64559.peg.2285 | gene-AT00_RS12560 | Putative 3-oxoacyl-[ACP] synthase |
| APE_N | 6666666.166623.peg.888 | 6666666.64559.peg.2286 | gene-AT00_RS12565 | Conserved hypothetical protein |
| APE_M | 6666666.166623.peg.890 | 6666666.64559.peg.2287 | gene-AT00_RS12575 | Conserved hypothetical protein |
| APE_L |  | 6666666.64559.peg.2288 |  | Conserved hypothetical protein |
| APE_K | 6666666.166623.peg.892 | 6666666.64559.peg.2289 | gene-AT00_RS12585 | Hypothetical protein |
| APE_J | 6666666.166623.peg.894 | 6666666.64559.peg.2291 | gene-AT00_RS12600 | Putative enzyme |
| APE_J | 6666666.166623.peg.895 |  |  | Putative enzyme |
| APE_I | 6666666.166623.peg.896 | 6666666.64559.peg.2292 | gene-AT00_RS12605 | Conserved hypothetical protein |
| APE_H | 6666666.166623.peg.897 | 6666666.64559.peg.2293 | gene-AT00_RS12610 | Putative enzyme |
| APE_G | 6666666.166623.peg.898 | 6666666.64559.peg.2294 | gene-AT00_RS12615 | Conserved hypothetical protein |
| APE_F | 6666666.166623.peg.899 | 6666666.64559.peg.2295 | gene-AT00_RS12620 | Putative acyl carrier protein |
| APE_E | 6666666.166623.peg.900 | 6666666.64559.peg.2296 | gene-AT00_RS12625 | Putative acyl carrier protein |
| APE_D | 6666666.166623.peg.901 | 6666666.64559.peg.2297 | gene-AT00_RS12630 | Putative phospholipid biosynthesis acyltransferase |
| APE_C | 6666666.166623.peg.902 | 6666666.64559.peg.2298 | gene-AT00_RS12635 | Conserved hypothetical protein |
| APE_A |  | 6666666.64559.peg.2300 |  | Putative O-methyltransferase |
| APE_B |  | 6666666.64559.peg.2299 |  | Conserved hypothetical protein |

**Supplemental File 1.**
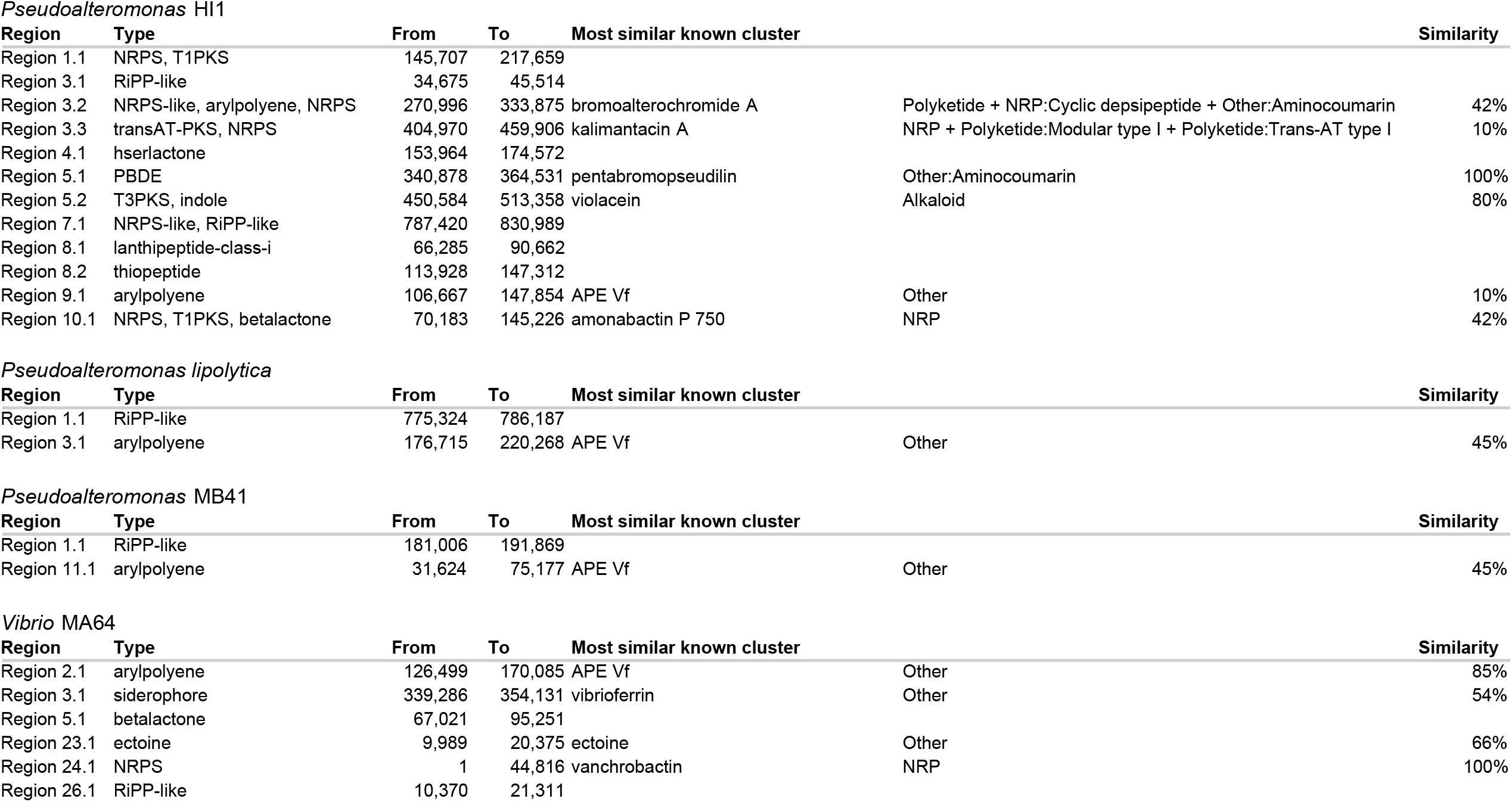
Genes shared between MB41 and MA64 identified with OrthoFinder.

